# A unified pipeline for discovering previously unknown enzyme activities

**DOI:** 10.64898/2026.02.02.703255

**Authors:** Ariane Mora, Julia C. Reisenbauer, Helen Schmid, Ikumi Miyazaki, Yueming Long, Jason Yang, Ryen O’Meara, Frances H. Arnold

**Affiliations:** Division of Chemistry and Chemical Engineering, California Institute of Technology, California Blvd., 91125, California, USA; AITHYRA GmbH Research Institute for Biomedical Artificial Intelligence of the Austrian Academy of Sciences Helmut-Qualtinger-Gasse 2, Stg. 2, 1030 Vienna, Austria; Institute of Science and Technology Austria (ISTA), Am Campus 1, 3400 Klosterneuburg, Austria; Department of Biosystems Science and Engineering, ETH Zürich, Basel, Switzerland; Department of Chemistry, Graduate School of Science, The University of Tokyo, Bunkyo-ku

## Abstract

Enzymes catalyze diverse chemical transformations and offer a sustainable approach to both breaking and making chemical bonds. However, finding an enzyme capable of performing a specific chemical reaction remains a challenge. We developed a new framework, Enzyme-toolkit (Enzyme-tk), that integrates 23 open-source tools to enable the discovery of enzymes that have activity toward a specific target reaction. Additionally, we introduce two new methods to facilitate enzyme discovery: (1) Func-e, an ML tool that searches large databases for enzymes that potentially catalyze a specific chemical transformation and (2) Oligopoolio, a gene assembly approach that reduces the cost of accessing protein sequences and thus the barrier to their experimental validation. We applied Enzyme-tk to find enzymes for chemical degradation of two man-made pollutants, di-(2-ethylhexyl) phthalate (DEHP) and triphenyl phosphate (TPP). We demonstrate that new, previously unannotated enzymes with favorable characteristics, such as high thermostability, can be identified using Enzyme-tk for reactions that are dissimilar to the training set.

## Introduction

Enzymes can catalyze a plethora of chemical transformations and are used across many industries, from pharmaceutical manufacturing to degradation of environmental pollutants. However, finding an enzyme that performs a specific biocatalytic transformation remains a challenge, especially for unnatural substrates.^1^ These enzymatic starting points are usually discovered by screening arrays of diverse enzymes for a desired function, a resource- and time-intensive process. Popular webtools such as MetaCyc,^2^ KEGG,^3^ and others^4–6^ enable enzymes to be queried based on similarity. Results are then based on correlations between ligands or between native enzymatic reactions and the desired chemical transformation given by the user. Alternatively, protein databases such as BRENDA^7^ and UniProt^8^ can be mined to find enzymes that catalyze a specific known reaction or class of reactions. However, this discovery process remains *ad hoc* and non-standardized, requiring users to navigate across many resources, either manually filtering or writing bespoke scripts to select for specific characteristics.

Recently, machine learning (ML) methods have been presented to complement sequence- and substrate-based approaches to discover enzymes with no annotated homologs or similar (natural) reactions.^9^ Several tools have been developed recently to assign a reaction or an enzyme to an enzyme commission (EC) number or the enzyme’s ability to perform a given reaction.^10–12^ As with similarity searching tools, there is no standard format for files or the environment in which to run these tools. Finally, most ML methods do not experimentally validate predictions, especially for novel enzymatic functions on non-natural substrates.^10,12,13^

For new or unannotated chemical transformations, both similarity and ML search methods have shown limited generalizability to “out-of-distribution” reactions for novel transformations or substrates that are currently not captured in databases.^13^ Experimental success rates for finding active enzymes for even annotated functions remain low and significantly drop for novel transformations.^14^ In most cases, prediction methods are only tested *in silico*,^15^ in part owing to the high cost of experimental validation and lack of standardized methods for closing the ML-experimental loop. The cost of a full-length gene resulting in a 200 amino acid (AA) sequence is between $70 and $170 (Twist, IDT, September 2025). Several recent approaches have proposed new methods leveraging pools of oligomers that consist of multiple gene fragments, assembling full-length genes using Golden Gate assembly^16–18^ or using Polymerase Cycling Assembly (PCA).^19–21^ There currently exists no standard pipeline to discover biocatalysts, with tools fragmented between the chemistry and the protein domains, and no guidelines for the integration of experimental data.

To fill this gap, we introduce a framework for enzyme discovery and engineering from diverse data sources, beginning with ML discovery and including both the experimental workflow and feedback loop. Enzyme-tk includes 23 open-source tools, in addition to a new ML classifier, Func-e, to discover enzymes across large databases that may perform specific reactions and an experimental protocol to enable the evaluation of enzymes assembled via a cost-effective gene assembly method, Oligopoolio. We applied our approach to two candidate reactions with unnatural substrates, finding enzymes that catalyze the first step of the degradation pathways of commonly found pollutants. We envision that this work will serve as a core framework for ML engineers who may aim to improve and add new open-source ML models and experimentalists who wish to use the tools to facilitate experimental workflows for discovering new enzymes for further engineering.

## Results

### A generalizable pipeline for enzyme discovery

We developed a new pipeline to bridge the computational and experimental gap in ML-driven discovery of enzymes. The Enzyme-tk pipeline has three core modules: ‘predict’, ‘synthesize’, and ‘validate’, Figure 1.A. Enzyme-tk is a Python package consisting of 23 open-source tools that are composable. Enzyme-tk enables researchers to turn a large set of unannotated sequence reads and a chemical transformation into filtered designs of gene constructs for testing enzymatic activities, Figure 1.B. Standardized data formats are passed between tools to bypass conversion between FASTA, CSV, and other output file formats, see Table 1 for a list of tools. Each module ingests and outputs CSV data, making visualization and interaction simple for researchers. Furthermore, Enzyme-tk handles the different computational environments for each module by using conda, a python environment manager^22^, automatically activating and deactivating respective conda environments. Enzyme-tk is designed with modularity as a principle, which facilitates different enzyme discovery objectives and is designed to be easily extensible by the broader enzyme community. Figure 1.B provides an overview of all modules of the pipeline which can be interchangeably run using a single line of code.

**Table 1:**
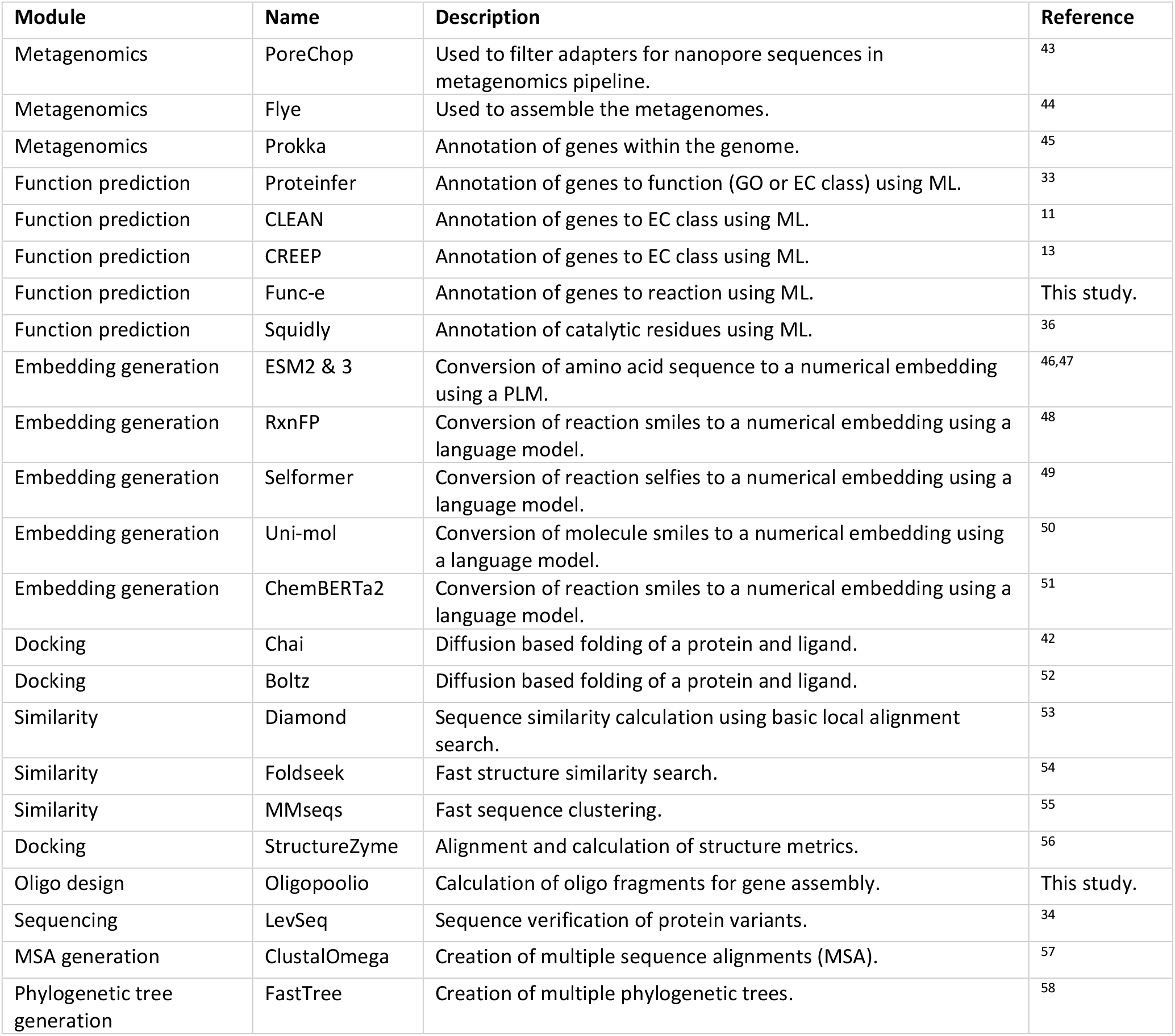
Description of tools currently available in Enzyme-tk. Some tools contain multiple functions, however, the main function used in Enzyme-tk has been listed in the table.

**Figure 1.**
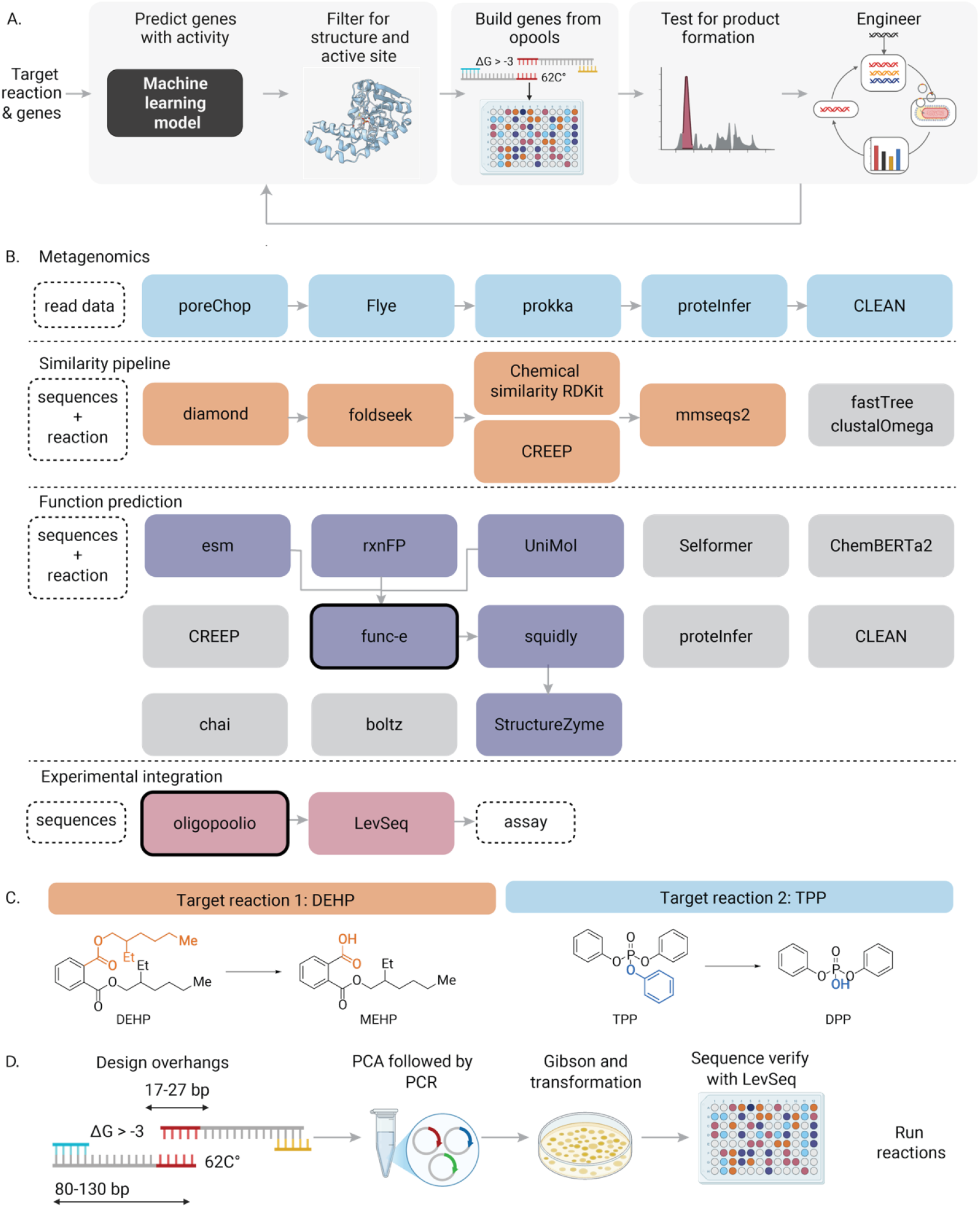
A. High-level overview of the Enzyme-tk pipeline. Users input a desired target reaction and a sequence database on which they wish to predict activity for, the program then predicts which sequences within this set are most likely to be able to catalyze the reaction. Predictions are filtered using structural and active site predictions. Genes are then synthesized using an oligomer assembly method, Oligopoolio, designed to reduce the cost of gene synthesis from oligopools (opools). Experimental validation is performed by testing for a specific target transformation. Successful variants can be further explored and engineered for a desired process or re-included in the ML loop, depending on the scientific or engineering objective. B. 23 computational open-source tools are currently included in Enzyme-tk, here organized around three pipelines created for this study, which are shown highlighted with the colored boxes. Each module can be linked together interchangeably or substituted by alternative tools (highlighted in grey boxes; e.g. in the function prediction module, CLEAN could be used instead of Func-e). Novel tools introduced in this study (black outlined boxes). C. Target reactions of the two partial degradation pathways, 1) hydrolysis of di-(2-ethylhexyl) phthalate (DEHP) to mono(2-ethylhexyl) phthalate (MEHP) and 2) hydrolysis of triphenyl phosphate (TPP) to diphenyl phosphate (DPP). D. Overview of the Oligopoolio pipeline: overhangs are automatically designed, ordered from a DNA provider, and assembled with a polymerase cycling assembly step, followed by PCR to amplify the full assemblies. Circular vectors are created using Gibson assembly and transformed into E. coli. Inserts are sequence-verified for accuracy with LevSeq^34^.

To validate the utility of this pipeline, we sought to identify enzymes capable of performing two specific degradation reactions, 1) hydrolysis of di-(2-ethylhexyl) phthalate (DEHP) to mono(2-ethylhexyl) phthalate (MEHP) and 2) hydrolysis of triphenyl phosphate (TPP) to diphenyl phosphate (DPP), Figure 1.C. DEHP and TPP are representatives of two classes of common pollutants, phthalates and organophosphates. Phthalates are commonly used as plasticizers and, owing to their high use and bioaccumulation, are found abundantly in surface waters and wastewater biosolids.^23^ The higher molecular weight species, such as endocrine-disrupting DEHP,^24^ pose challenges for natural degradation owing to their sterically demanding side chains.^25^ Similar degradation challenges have been encountered for some organophosphate esters, such as TPP, which are used as flame retardants, industrial additives, and plasticizers.^26,27^ This phosphate ester has also been linked to reproductive and aquatic toxicity.^28,29,30^

The enzymes already reported to perform these target transformations are not ideal candidates for real-world applications, such as in wastewater plants. Their large size (QHH21706.1 is an esterase from *Bacillus subtilis*^31^ that comprises 489 amino acids (AA) and *Sb-*PTE is a phosphotriesterase from Sphingobium sp. TCM1 comprising 583 AA)^26^ and structural features (*Sb-*PTE contains disulfide bonds) limit their utility. Given the limitations of these existing enzymes, herein referred to as QHH (ID: 5387) and *Sb-*PTE (ID: 5388), we sought to find alternative enzymes with high activity and more favorable characteristics (small, thermostable, and high expression levels in *Escherichia coli*).

To search for potential enzymes, we curated two discovery sets, an “extremophile dataset” and a “metagenomics dataset.” The first dataset consists of 8,793 enzymes from UniProt with favorable characteristics: short sequences (100–350 AAs) from bacterial or archaeal thermostable organisms (>50 °C).^32^ For the second dataset, we sought to investigate mining unannotated microbial sequences and selected a metagenomics dataset originating from the influent and activated sewage sludge from five wastewater treatment plants across India, the United States, Switzerland, Sweden, and Hong Kong (see Methods for processing details). We filtered the dataset to 2,708 putative esterases by predicting the EC numbers using two ML tools, proteInfer^33^ and CLEAN^11^, see Methods for details.

### Experimental validation of baseline computational methods

To test the discovery loop using conventional methods, we applied two existing approaches to identify potential enzymes. The first is a baseline approach in which sequences are selected based on substrate similarity, and the second is our previously published multi-modal ML approach, CREEP, that uses contrastive learning to embed sequences into a joint space, enabling sequences to be extracted based on reactions that are projected “close” in the latent space.^13^ Reviewed bacterial or archaeal sequences from SwissProt were filtered for the following EC classes: DEHP: 3.1.1.74, 3.1.1.1, 3.1.1.3; TPP: 3.1.3.48, 3.1.4.4, 3.1.3.2, 1.7.1.6; CREEP ML embedding approach based on DEHP: 3.1.1.3, 3.1.1.101, 3.1.1.74, 3.5.2.6, 3.1.2.2, 3.3.2.10, 3.1.1.76.), resulting in 40 putative enzymes to test for activity, see Methods for details.

The high cost of gene synthesis motivated us to design a new approach to build genes from fragments using pooled Polymerase Cycling Assembly (PCA), Figure 1.D. Ordering gene fragments in a pooled fashion reduces the cost of gene synthesis for a 96-well plate with IDT by 45%. Using the Oligopoolio submodule of the pipeline to select the optimal fragment size and overlaps, 40 genes were ordered with fragments ranging from 414 to 894 base pairs to test the assembly method. A two-step PCR assembly protocol was developed, see Methods. Full-length genes were identified and sequence-verified using LevSeq^34^ in 96-well plate format. We found that 34/40 genes were correctly assembled with 100% sequence identity (> 3.5 pmol/oligo/μL, IDT oligomer pool). Shorter genes assembled more efficiently. Hence, to ensure representation of longer sequences, at least five clones per variant needed to be picked, see SI Section 3 for a comparison to existing methods, S. Table 13 and assembly statistics, S. Figure 2-8. S. Table, 14-16.

The assembled genes were overexpressed in *E. coli* and screened in 96-well plate format for the degradation of DEHP and TPP using clarified cell lysate (see Methods and SI for details). To efficiently detect product formation in our experimental pipeline, we employed relatively high substrate loadings (DEHP: 3.9 g/L; TPP: 326 mg/L), which greatly exceed concentrations typically found in wastewater streams (e.g. DEHP: 0–8840 ng/L^25^; TPP: 0–491 ng/L^35^). Although no enzymes had activity comparable to the enzymes known to have these activities (positive controls), a low level of activity was observed for the carboxylesterase Q06174 (ID: 5395) on TPP, S. Figure 22. To demonstrate the ability to select candidates with desirable traits such as engineerability, we performed one round of random mutagenesis (*ep*PCR) on Q06174, see SI Section 2 and 6 for details. We identified five variants that exhibited increased activity compared to Q06174 under validation conditions. These mutants exhibited activity on TPP equivalent to the positive control, with the best Q06174-3 variant harboring the K138E, D178G, and D236G mutations (ID: 5396), Figure 2. No enzymes were identified with activity for DEHP hydrolysis.

**Figure 2.**
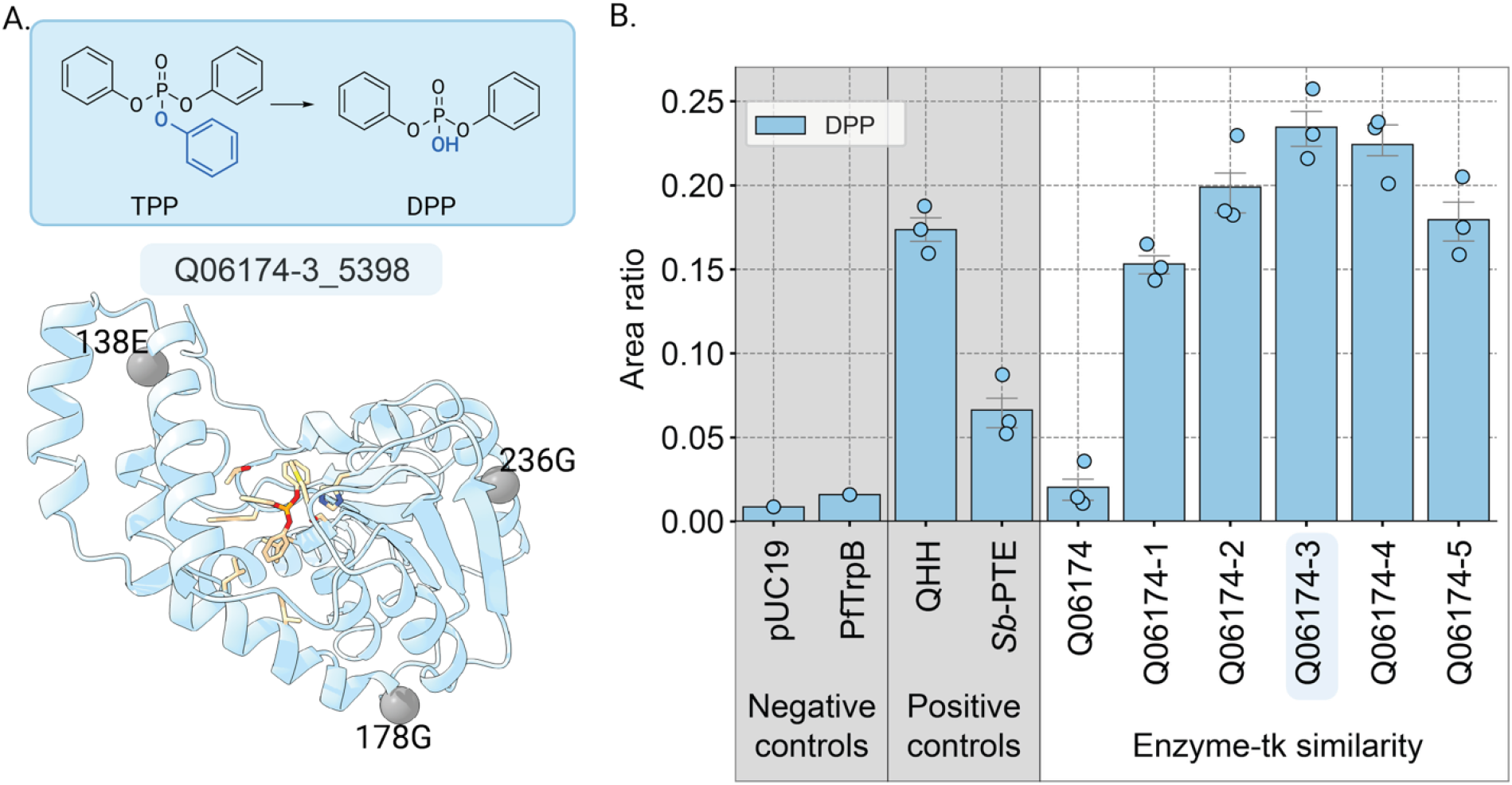
A. Structure of Q06174 co-folded with TPP using Chai^42^ showing the substitutions of the best performing variant Q06174-3 (with K138E_D178G_D236G substitutions). B. Validation results for the epPCR library showing area ratio (integrated product area/integrated internal standard area) for the formation of DPP from TPP. Reactions were performed in technical triplicate under validation conditions, and papaverine hydrochloride was used as an internal standard when analyzing product formation via HPLC-(MS)-analysis.

### Improving performance with a new ML method for searching sequence space

Given the limited success of the EC-based approaches for identifying enzymes capable of the hydrolysis of DEHP and TPP, we developed a new sequence-only based ML model, Func-e, to predict the activity of an enzyme for a desired reaction. The model uses embeddings of the enzyme sequence and chemical reaction space and combines these with two cross attention heads. These are then fed into a neural network with a prediction head and 13 regression heads that predict features such as molecular weight and sequence length (see Figure 3.A for model architecture). The intersection of the reaction and protein subsets from CARE^13^, a recently published benchmark to measure the performance of enzyme function prediction tools, was used for training and *in silico* evaluations (see Methods for details, S. Table 1 for dataset sizes, S. Table 2 for similarity to target reactions and positive controls). After training for 20 epochs, the model shows some *in-silico* generalization capacity, with an expected drop in performance as the test set becomes more dissimilar to the training set, S. Figure 1. The model that was used to inform the final architecture had an accuracy of 0.75, 0.61, and 0.56 on a balanced dataset with sequences which were less than 50% similar to the training set, on the easy (held out reactions from EC 4), medium (entire held out classes from EC 4), and hard test sets (entire held out classes from EC 3 classes), see SI Section1 for details.

**Figure 3.**
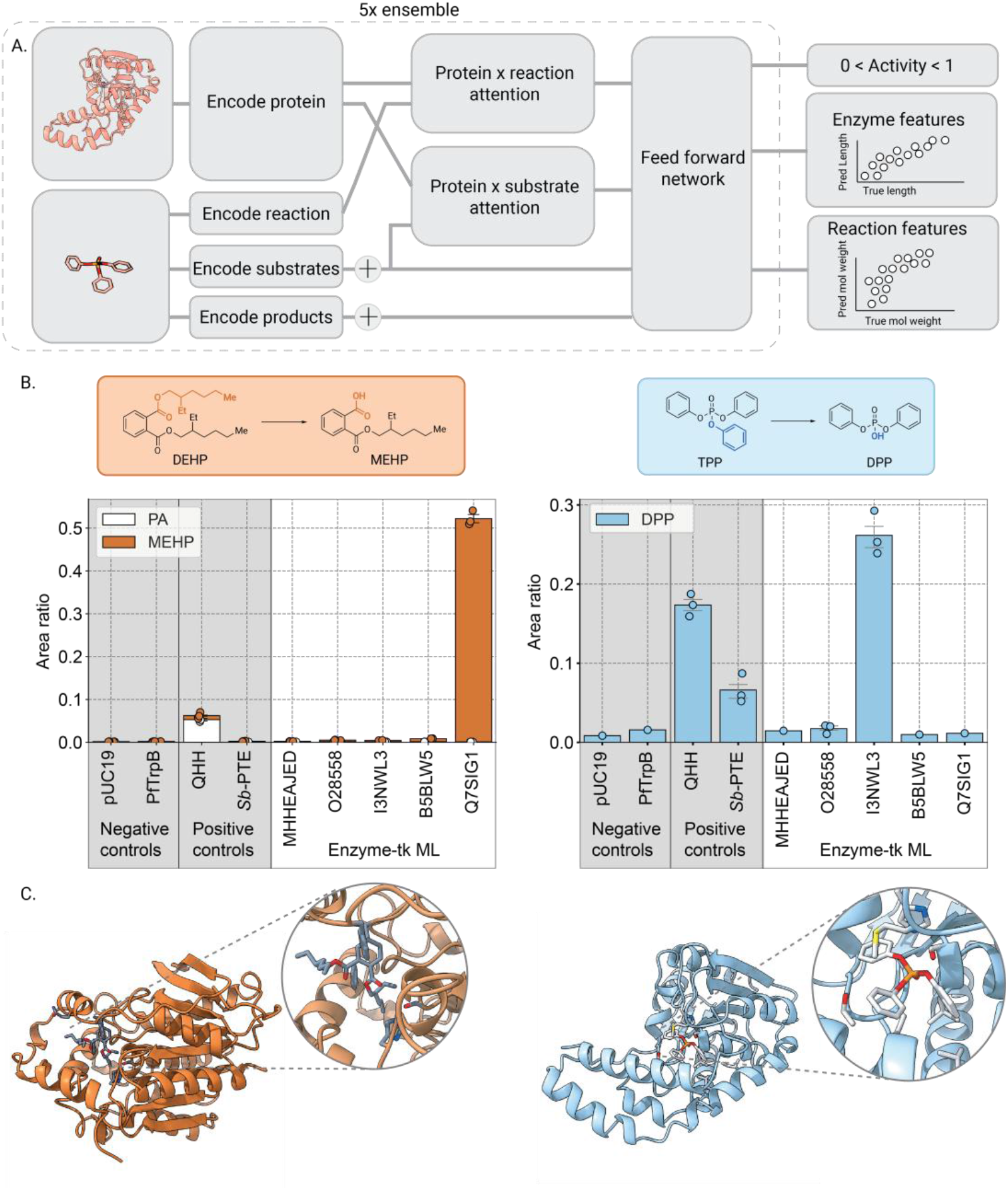
A. Overview of the architecture of our predictive ML model, Func-e. B. Validation to test enzyme activity towards MEHP and DPP formation. Clarified cell lysate reactions were performed under validation conditions in technical triplicate, and papaverine hydrochloride was used as an internal standard when analyzed by HPLC-(MS)-analysis. Results show integrated product area normalized by the internal standard area for each enzyme and target reaction. PUC19 and PfTrpB are negative controls, QHH is a positive control for DEHP to PA hydrolysis and additionally has promiscuous activity towards TPP degradation. Sb-PTE is a positive control for the TPP to DPP hydrolysis. C. Co-folded structure using Chai for Q7SIG1 with the DEHP substrate and I3NWL3 with the TPP substrate.

In addition to applying the model to the metagenomics set described above, we increased the search space of the extremophile dataset to include unreviewed enzymes from UniProt, resulting in a dataset size of 8,793 putative enzymes. Enzymes with a mean prediction >0.5 and mean distance between the predicted length and the true sequence length of < 50 amino acids were selected. The predicted enzymes were passed through our previously developed active site annotation filter^36^ and structural filtering pipeline (https://github.com/moragroup/StructureZyme, see Methods for details). Based on manual inspection of the structures, 14 enzymes from the extremophiles and 17 from the metagenomics set were selected for experimental validation. The 31 predicted genes were assembled using Oligopoolio and screened in 96-well plate format, S.Table 17 for plate format and SI Section 5 for experimental results. Five enzymes showed potential activity and were selected for validation, four from the extremophile dataset and one metagenomics sequence, S.Figure 12 and 16. Of these, two showed significantly higher activity toward the desired reaction compared to the best enzymes from the literature, Q7SIG1 (ID: 5389) for DEHP and I3NWL3 (ID: 5391) for TPP hydrolysis, Figure 3B-C.

## Discussion

A significant problem in enzyme engineering is how to best discover an enzyme with activity towards a specific reaction. Recent reports have sought to enhance the identification of new enzymes by using ML to annotate uncharacterized sequences to EC numbers in low homology regimes^11^ and new reactions to EC numbers.^13^ Building on this work, we developed a ML model, Func-e, to predict whether an enzyme can perform a specific reaction. In combination with this core ML module, we consolidated over twenty publicly available tools and unified them in a single pipeline, Enzyme-tk, to streamline the workflow to identify suitable enzymatic starting points with desirable traits for a specific catalytic function from a user-specified “sequence search space”. In addition to the commonly used sequences in SwissProt, datasets were curated to increase the sequence search space for potentially effective biocatalysts. Finally, to reduce the cost of the experimental pipeline, a gene assembly method based on oligos, Oligopoolio, was developed as a sub-module to bridge the gap between *in silico* and experimental testing.

Oliogopoolio is a new joint computational and experimental method for assembling full-length genes using low-cost oligo pools. While various approaches have been reported for achieving similar tasks,^16,17,20,37–39^ efficient command line tools for the generation of constructs from multiple distinct sequences using PCA in a single test tube remain scarce. Oligopoolio was used to assemble multiple pools with varying success, often reaching 100% recovery of all desired genes in a single oligo pool sample. However, we noticed that a higher concentration of DNA in the oligo pool drastically impacted the probability of producing sequence-correct variants. Hence, we suggest using 50 pmol pools from IDT (> 3.5 pmol/oligo/μL, IDT opool). With further methodological optimization, we believe that the protocol could also be amenable to lower concentrations of oligo pools. This module of the pipeline could also be swapped for an alternative method such as Golden Gate assembly^16^. Not designed for highly similar genes or for large gene assemblies, Oligopoolio instead bridges the gap between ordering single genes and a plate of diverse genes in a cheap and accessible manner. Assembly of highly similar genes would likely result in chimeric products, and the success rate of assembly in larger pools could be limited by the DNA concentration. We find the oligo pool approach to be an effective method for not only reducing the cost of synthesis but also minimizing the use of expensive enzyme stocks and reagents, as it requires only two pooled PCRs to construct all desired genes in a single tube within a single day.

With the modular Enzyme-tk pipeline in hand, we first evaluated the performance of a similarity-based module, instead of the more complex Func-e ML tool, to filter for potential enzyme candidates applied to SwissProt sequences. This subset of the pipeline enabled us to identify an enzyme, Q06174, from an extremophilic thermostable organism catalyzing the hydrolysis of TPP to DPP. One additional round of directed evolution provided Q06174-3 (with K138E_D178G_D236G substitutions; ID: 5396), which is less than half the length of literature-known *Sb*-PTE and exhibits modest activity toward TPP hydrolysis. This demonstrates the modularity and compact workflow from a sequence dataset to filtering and validation of capable enzyme variants.

To further extend the search space to our curated metagenomics and extremophile datasets, the developed ML model, Func-e, was implemented to search for enzymes for the partial degradation of DEHP into MEHP and TPP into DPP. Neither of the tested reaction smiles are similar to the training set, for DEHP, the closest reaction similarity is 0.46 and for TPP it is 0.48, using Tanimoto similarity with structural fingerprints, S. Table 1. *In-silico* predicted enzymes were then assembled via Oligopoolio and experimentally evaluated. For the DEHP hydrolysis, an unreviewed enzyme from Uniprot, Q7SIG1 (ID: 5389), originating from thermophilic *Bacillus acidocaldarius*, demonstrated catalytic activity. Q7SIG1 is a hydrolase (no EC annotation in UniProt^40^), which is able to catalyze the first hydrolysis step in DEHP degradation. This is in contrast to the literature enzyme QHH, which is able to fully degrade DEHP into phthalic acid (PA). Though this is unsurprising, as the first degradation step of this transformation was the sole input into the ML model. For the partial TPP degradation, another unreviewed carboxylesterase originating from *Geobacillus sp*. ZH1, I3NWL3 (ID: 5391, EC:3.1.1.1, 246 AA length), was identified, which is 337 AAs shorter than previously reported *Sb-*PTE. Translating to a less than 30% sequence identity between the newly discovered enzymes and the positive control sequences,^41^ demonstrating the predictive and filtering capabilities of our computational framework. Neither of these enzymes would have been discovered using a simple EC search on annotated data: I3NWL3 was active on TPP, yet EC3.1.1.1 was not predicted as a top EC for this reaction, while for Q7SIG1 no EC number is assigned in UniProt. Both enzymes were identified by screening fewer than 30 enzymes (out of 8,793 thermophilic enzymes considered), reducing experimental cost and discovery time. Of note, given the high lipophilicity of these substrates and the associated risk of phase separation at such concentrations under aqueous conditions, a qualitative comparison is most appropriate under the developed analytical conditions and sufficient for finding initial activities that can be improved by protein engineering/directed evolution.

In contrast to other models,^10^ we opted to design a protein sequence-only method to enable rapid *in-silico* filtering of predictions, before passing the predicted sequences to more computationally intensive structural filtering steps. In future work, ensembling of multiple ML models could potentially improve the activity prediction, further increasing the success rate of finding active enzymes within a small pool of experimentally tested variants. The pipeline has been designed modularly to enable exactly such integrations.

## Conclusion

We designed Enzyme-tk to be a platform for the prediction, synthesis, and experimental validation of enzymes with a desired catalytic activity. The code is fully open source to enable the composition of sub-modules. This modularity will allow its adoption across disciplines, where enzymes with specific traits for desired chemical transformations need to be identified or improved. In showcasing the utility of the pipeline for two bioremediation tasks, we also provide the community multiple ready-to-use pipelines that will be of broad utility, including 1) metagenomics reads to EC classification and reaction prediction, 2) sequence filtering for enzyme traits, prediction, active site annotation, structural filtering, and oligo design. These pipelines along with documentation are available on our GitHub (https://github.com/ArianeMora/enzyme-tk). We hope to build a thriving open-source community that intersects the enzyme annotation and enzyme design domains.

## Author contributions

A.M., J.C.R., and F.H.A. conceived the idea. A.M. developed the ML model. A.M., H.S., and I.M. developed the computational pipeline, with input from R.L.O. on the dataset conceptualization and J.C.R., J.Y., and Y.L. on the pipeline conceptualization. H.S. developed the structural pipeline. J.Y. ran the CREEP model. J.C.R. developed screening assays and the experimental testing pipeline. J.C.R., H.S., I.M., A.M., R.L.O., and Y.L. performed experimental validations. A.M., J.C.R., H.S., and I.M. developed the oligopool assembly strategy. A.M. and Y.L. developed the oligopool sequencing analysis pipeline. A.M., J.C.R., and F.H.A. wrote the manuscript, and all authors edited and revised the manuscript.

## Acknowledgments

The authors would like to thank Sabine Brinkmann-Chen for careful proofreading and valuable discussions, and Jarrid Rector-Brooks for careful proofreading.

## Materials and Methods

### Data selection and processing

Data were downloaded from UniProt on the 17^th^ of February 2025 by selecting reviewed sequences and the following additional information (Fragment, Length, Mass, Organism ID, Taxonomic Lineage (IDs), Active Site, Cofactor, EC number, Reviewed), totaling 572,970 protein sequences. These were filtered for size (minimum length 100, maximum 1024), to be non-fragments, and proteins with an EC annotation, resulting in 261,254 potential enzymes. For each sequence, the mean summarized esm3 embeddings (esm3-open, mean of the per-token-embedding)^46,47^ were computed.

Reaction data were downloaded from EnzymeMap,^59^ containing 62,896 deduplicated reactions, or 54,090 products and 37,263 substrates. Properties were calculated for each substrate and product by using RdKit version 2024.3.3, calculated properties were molecular weight (MolWt), total polar surface area (TPSA), calculated solubility proxy (MolLogP), and the maximum and minimum Gasteiger partial charges (MaxPartialCharge and MinPartialCharge). For reactions with multiple substrates and products, these were summed to create a single embedding. For each reaction, chemberta2^51^ and rxnFP^48^, additionally for each substrate and product the uniMol^50^ representation was calculated.

### Metagenomics data processing pipeline

Long-read metagenomics data were downloaded from a study^60^ that investigated antibiotic resistance genes both from influent and activated sewerage sludge from five wastewater treatment plants from India, the United States, Switzerland, Sweden, and Hong Kong. These were available via the SRA under the following accessions, SRX8190902, SRX8190901, SRX8190900, SRX8190899, SRX8190898, SRX8190897. SRA toolkit was used to download and unzip the data. Porechop^43^ was used to trim the nanopore adaptors, before Flye was used to assemble the metagenomes.^44^ Genes were annotated to the genomes using Prokka,^45^ before enzyme commission numbers were predicted for each gene using ProteInfer^33^ and CLEAN.^11^ Default parameters were used for all tools. Genes were retained if they fell within the EC numbers, EC:3.1 or 3.- (i.e. retained if unable to be assigned at the second level), and finally, filtered for length (100 < length < 900 amino acids). These genes were used as the “metagenomics set” referred to herein.

### Extremophile dataset processing

The temperature was annotated to the enzymes using data from^32^ a resource containing the optimal growth temperature data for 21,498 organisms. Of the 261,254 UniProt enzymes, 218,840 were able to be mapped to an organism temperature. Growth temperature is used as a proxy for thermostability. Temperature annotations were mapped at the species level if it existed, otherwise genus, then family, and order. If no order was identified, temperature was omitted from this sample. After the first pass on the EC classification dataset, we extended this to unreviewed sequences in UniProt (filtered for proteins with existence at 1-2, or annotation level of 4 or 5), resulting in a dataset of 8,793 thermophilic enzymes.

### Literature search

To find a positive control for the degradation of DEHP and TPP, scientific literature was searched by using the terms “DEHP/TPP enzymatic degradation”, finding several papers that met our criteria. The final selection was based on reports describing the culturing, isolation, and testing of specific enzymes in detail, selecting QHH21706.1, a *para*-nitrobenzyl esterase from *Bacillus subtilis*,^31^ and *Sb-*PTE, a phosphotriesterase from *Sphingobium sp*. TCM1, as our positive controls.^26^

### Training and test sets setup

Training and test sets were created using the splits as defined in CARE.^13^ The reaction and enzyme datasets were combined by taking the intersection of the training and test sets for protein similarity 30-50 (EC’s and enzymes). These sets were filtered to ensure no contamination between the test and training sets, by running diamondBlast^53^(v 2.1.11) against the training set for each test set (sequence similarity), and FoldSeek^54^(v 10.941cd33) (structural similarity). The PDB was used as the database in FoldSeek with the pdb structures mapped to their respective UniProt identifier. Any training data entries that mapped to the test sets with an identity > 0.5 (both structural and sequence) were omitted from the training datasets. The reactions for each EC were also tested for similarity to the training set, here, the reaction distance (Tanimoto similarity) from RDkit was used (v 2024.3.3). Note, some reactions had a high identity to a reaction in another EC class; in these instances, the reaction was considered ambiguous and removed from both EC classes.

To create the training and test sets, pairs of positive and negative data were created by selecting one reaction as a positive from the EC annotated to the enzyme and one as a negative from an EC number that was not annotated to the enzyme, and these were annotated as 1 and 0, respectively. This was done for each of the 184,529 enzymes in the training set for the protein space. Note only ECs from each of the reaction sets were used, making an “easy”, “medium”, and “hard” training set for reactions and enzymes. The test sets were created in a similar manner, where a sequence was annotated as positive for a single reaction for the annotated EC number and also assigned to another random EC as a negative data point, for which a complementary positive and negative data point was also created. Different EC number levels were used to make easier and more challenging training points.

### ML Model, func-e

We designed a new ML model, func-e, with a binary class classifier with 13 regression heads, with five enzyme features: enzyme length, mass, polarity, temperature, and the total polar surface area, and four features each from the substrates and the products: molecular weight (MolWt), total polar surface area (TPSA), calculated solubility proxy (MolLogP), and the maximum and minimum Gasteiger partial charges (MaxPartialCharge and MinPartialCharge). The model had a self-attention mechanism on the substrate (uniMol), product (uniMol), reaction (rxnFP), and enzyme (esm) heads. This was concatenated with a bidirectional cross attention head (eight heads, 128 embedding size) between the substrate and the enzyme, and the reaction and the enzyme. The attention was added to the unaltered inputs for substrate (N=768) and enzyme (N=1280), before being fed through a feedforward neural network of three layers, 1024, 512, and 256. The loss was calculated as the mean of activity (binary cross entropy loss), and the masked mean-squared error (MSE) of the regression features. Dropout layers were used (0.2), with a 1,000 batch size, learning rate of 0.01 and with early stopping for a maximum of 30 epochs. An ensemble was created across five models with the mean prediction and variance recorded across each model. The ensemble was selected across four EC classes with different inferred negatives selected for each, and the final models were trained using the “easy” training set, for further details see SI Section 1. The reaction used as input was the smiles string: CCCCC(COC(C1=CC=CC=C1C(OCC(CCCC)CC)=O)=O)CC>>CCCCC(COC(C2=CC=CC=C2C(O)=O)=O)CC for DEHP and O=P(OC1=CC=CC=C1)(OC2=CC=CC=C2)OC3=CC=CC=C3>>O=P(O)(OC4=CC=CC=C4)OC5=CC=CC=C5 for TPP.

### Structural pipeline

Final candidate selections for experimental validations were performed after manual evaluation of all filtered enzymes using a structural pipeline (https://github.com/moragroup/StructureZyme). In short, enzymes were first assigned catalytic residues using our recently developed catalytic residue prediction model, Squidly.^36^ Structures were then co-folded with the predicted substrate (TPP or DEHP) using Chai^42^ and Boltz2^52^. If Alpha Fold 2 (AF2)^61^ structures were in the AF2 database, then these were also collected. The physics-based tool autodock Vina^62^ was used to then re-dock the substrate in either the AF2 or Chai structure. Metrics from each of the structural tools were extracted. Additionally, for each protein, pairwise superimposition over the predicted structures was performed, and features were extracted using PLIP^63^. The distance between the ligand and the nucleophile was calculated using the docked structures in Python. Enzymes were also manually visualized and selected for further validation.

### Annotation filtering steps based on UniProt

Initially, we aimed to keep our filtering to a minimum, filtering enzymes for origin (bacteria and archaea), length (<300 AA), simple structures (monomers/homodimers), and for the hydrolysis enzymes, no cofactors (i.e. ester cleavage by a catalytic triad). In the second step, we reduced the constraints of no cofactors and instead added an extra constraint of expression in the cytoplasm.

### CREEP Model baseline

We trained a new CREEP model, using all available (no held-out) reaction-protein pairwise data, as given by the reaction2EC.csv and protein2EC.csv files in the CREEP repository. Afterward, we performed retrieval across all the SwissProt enzymes compiled in CREEP (protein2EC.csv) by searching based on the first step in the DEHP degradation. The top 1000 retrieved enzyme sequences (not grouped by EC number) were then filtered to include only short (<350 AA) archaeal or bacterial sequences, that were not annotated to require a cofactor and formed either a homodimer or monomer leaving 25 sequences. Of these, we removed EC numbers outside the class of hydrolases, selecting the following EC classes, 3.1.1.3, 3.1.1.101, 3.1.1.74, 3.5.2.6, 3.1.1.-, 3.1.1.3, 3.1.2.2, 3.3.2.10, and 3.1.1.76.

### Similarity selection baseline

In addition to enzymes selected based on machine learning, we sought to benchmark these compared with a similarity-based approach, searching the reviewed protein space in UniProt. We tested the baseline substrate similarity approach by searching for proteins that use the most similar substrates to a target reaction, using the Tanimoto similarity using Morgan Fingerprints with parameters from Rdkit (radius=2, fpSize=2048). For TPP and DEHP, we filter based on sequence length (<300 AA), origin (bacteria or archaea), cofactors (none), and oligomeric state filter (homodimer or monomer). For TPP, this reduced the number of enzymes to 50. To remove redundancy, mmseqs2 was used to cluster sequences at 90% identity. From the 18 clusters, the centroid sequence from each cluster was selected. For DEHP, the initial filter left 12 enzymes to test. For the second set of filters, we also selected for cytosolic expression as some of the secreted enzymes may have disulfide bonds.

### Computational oligo assembly method (Oligopoolio)

Oligo-sequences are short sequences (<300 nt) that are generally less expensive per base pair compared to ordering full-length genes. We designed a computational approach using polymerase cycling assembly (PCA) to assemble full gene constructs from short oligos. The tool includes two computational choices, first, a swarm-based optimization algorithm that aims to optimize on multiple constraints using Pyswarms (v. 1.3.0), second a deterministic method that iteratively selects the optimal overhangs. The constraints are, melting temperature of 62 °C for overhangs, DG of homodimer > –3 kcal mol^-1^, not starting or terminating on a triplet (e.g. AAA), and having a minimum Hamming distance of eight to other overlaps. These are optimized by altering the cut sites along a gene, with the oligo lengths between 80 and 130 nts and the overlap or primer length between 18 and 27 nts. For the stochastic approach, as there are no guarantees as to reaching the global optimum, we perform the swarm five times and retain the oligo cut sites from the best cost. Each of these oligos is then iteratively flipped, starting with a reverse strand and ending either on a forward strand. These sequences are then collectively evaluated for overlaps with other oligos in the pool, and any minimal Hamming distances are reported to the user, see https://github.com/ArianeMora/oligopoolio for details.

### Experimental oligo assembly method (Oligopoolio)

Oligopools (opools) were resuspended according to the suppliers’ recommendations and assembled via a two-step PCR protocol. The initial PCR was aimed at receiving full-length genes with annealing temperatures (T_annealing_ = 61.8 °C) based on the overlap regions, while the second one served as a clean-up and amplification protocol (see SI Section 3 for details). After clean-up, these gene inserts were then cloned into a pET22b(+)-vector between the *Nde*l and *Xho*I restriction sites using Gibson protocol in a single sample.^64^ Freshly transformed competent *E. coli* (BL21(DE3)) cells were plated on Luria broth (LB) agar plates containing 100 μg/mL carbenicillin, and stationary incubated at 37 °C overnight. Single colonies were picked into a 96-deep well plate containing LB_Amp_ (500 μL) to prepare starter cultures and incubated at 220 rpm for 12–16 hours at 37 °C. To confirm fully assembled sequences of desired genes, LevSeq^34^ was implemented, and cells were stored in glycerol stocks (25% glycerol) at -80 °C until used further. 96-well plates were re-arrayed to only contain desired genes, and starter cultures were prepared. Final variants were sequence-verified using Laragen Inc. (Culver City, CA), the DNA and protein sequences can be found in SI Section 7.

### Screening in 96-well plate format

LB_Amp_ starting cultures (50 μL) were used to inoculate TB_Amp_ pre-expression cultures (950 μL). These were then incubated at 37 °C for 2.5 hours at 220 rpm. After cooling on ice, protein expression was induced by the addition of β-D-1-thiogalactopyranoside (IPTG, 0.5 mM) and incubated at 22 °C for 16 – 21 h at 220 rpm. After harvesting, cells were frozen at -20 °C at least overnight before further use. Chemical lysis using 10 mM KP_i_ buffer (pH = 8) was performed, and clarified cell lysate was used to perform reactions in 96-well plates (for details see SI). Each well contained 10 mM KP_i_ buffer (pH = 8, 690 μL) and the respective substrate (10 μL of 1 M DEHP stock in DMSO, [DEHP]_final_ = 10 mM; 10 μL of 100 mM TPP stock in DMSO, [TPP]_final_ = 1 mM) and clarified cell lysate (300 μL). Reactions were performed under aerobic conditions, sealed and shaken at 220 rpm for 4 hours at 22 °C. Reactions were quenched by the addition of acetonitrile and analyzed by HPLC-(MS) analysis.

### Experimental validation

To validate identified hits from 96-well plate screens, glycerol stocks were streaked on LB_Carb_ agar plates and single colonies were picked to inoculate LB_Amp_ starter cultures (6 mL), which were used to inoculate TB_Amp_ pre-expression cultures (addition of 1 mL starter culture into 50 mL TB_Amp_). These were grown to an OD_500_ = 0.8 – 1.0, before cooling for 30 min on ice, and protein overexpression was induced with IPTG (0.5 mM). Cells were incubated at 22 °C for 16 – 21 h, then harvested by centrifugation, frozen overnight at – 20 °C, and chemical lysis was performed after thawing at room temperature by adding four times the volume of the wet-pellet weight (see SI Section 2 for procedures and SI Section 4 and 5 for results). Lysis buffer contained 10 mM KP_i_ buffer with 1 mg/mL lysozyme, 0.07 mg mL^-1^ DNAse, and 1μL mL^-1^ 2 M MgCl_2_. Clarified lysate was used to perform reactions in technical triplicate in 1.5-mL screw-cap glass vials. Each vial contained 10 mM KP_i_ buffer (pH = 8, 690 μL) and the respective substrate (10 μL of 1 M DEHP stock in DMSO, [DEHP]_final_ = 10 mM; 10 μL of 100 mM TPP stock in DMSO, [TPP]_final_ = 1 mM) and clarified lysate (300 μL). Protein concentrations were determined using the Pierce BCA Protein Assay Kit (Thermo Fisher Scientific) according to the manufacturer’s recommendations.

